# Comparative cytogenomics reveals genome reshuffling and centromere repositioning in the legume tribe Phaseoleae

**DOI:** 10.1101/2021.08.06.455448

**Authors:** Claudio Montenegro, Lívia do Vale Martins, Fernanda de Oliveira Bustamante, Ana Christina Brasileiro-Vidal, Andrea Pedrosa-Harand

## Abstract

The tribe Phaseoleae (Leguminosae; Papilionoideae) includes several legume crops with assembled genomes. Comparative genomic studies indicate the preservation of large genomic blocks among legumes, however, the chromosome dynamics during Phaseoleae evolution has not been investigated yet. We conducted a comparative genomic analysis to define an informative genomic block (GB) system and to reconstruct the ancestral Phaseoleae karyotype (APK). We defined the GBs based on the orthologous genes between *Phaseolus vulgaris* and *Vigna unguiculata* genomes. We searched for these GBs in different genome species belonging to the Phaseolinae (*P. lunatus*) and Glycininae subtribes (*Amphicarpaea edgeworthii* and *Spatholobus suberectus*), and in the *Medicago truncaluta* outgroup. To support our *in silico* analysis, we used oligo-FISH probes of *P. vulgaris* chromosomes 2 and 3 to paint the orthologous chromosomes of two non-sequenced Phaseolinae species (*Macroptilium atropurpureum* and *Lablab purpureus*). We inferred the APK with *n* = 11 and 19 GBs (A to S). We hypothesized five chromosome fusions that reduced the ancestral legume karyotype with *n* = 16 to *n* = 11 in APK. Furthermore, we identified the main rearrangements within Phaseolinae and observed an extensive centromere repositioning resulting from evolutionary new centromeres (ENC) in the *Phaseolus* lineage. Additionally, we demonstrated that the *A. edgeworthii* genome is more reshuffled than the dysploid *S. suberectus* genome, in which we could reconstruct the main events that lead the chromosome number reduction. The development of the GB system and the proposed APK provide useful approaches for future comparative genomic analyses of legume species.

## Introduction

Since the first plant genome sequencing and assembly (The Arabidopsis Genome Initiative 2000), genome sequencing technologies improved and became more accessible, increasing the genomic data available for economic and evolutionary important plant species (e.g., Li et al. 2018; Chen et al. 2018; Hu et al. 2021b). Sequencing genomes is essential for functional and comparative genomics, playing a fundamental role in understanding the plant biology and chromosomal evolutionary dynamics, such as genome reshuffling (Pavy et al. 2012; Cheng et al. 2014; Wang et al. 2021a), genome size variation (Wendel et al. 2016; Pellicer et al. 2018; Kreplak et al. 2019; Zhang et al. 2020a), polyploidy (Jiao et al. 2011; Soltis et al. 2015; Geiser et al. 2016; Ruprecht et al. 2017; Wu et al. 2020; Hu et al. 2021a), dysploidy (Lysak et al. 2006; Yang et al. 2014; Mandáková and Lysak 2018; Zhao et al. 2021), and other mechanisms for genome and species diversification.

Cytogenomic comparisons of related species provide important evolutionary insights. In Brassicaceae, for instance, chromosome painting based on *A. thaliana* BACs (Bacterial Artificial Chromosomes) as FISH (Fluorescent *in situ* Hybridization) probes (Lysak et al. 2001), together with a genomic block system (Schranz et al. 2006), elucidated the karyotype evolution within this family. These studies allowed to infer the ancestral crucifer karyotype (ACK), revealing chromosomal rearrangements related to the decreasing dysploidy in *A. thaliana* (Lysak et al. 2006), chromosomal reshuffling after whole genome triplication (WGT) in *Brassica* species (Cheng et al. 2014), and centromere repositioning across the family (Willing et al. 2015; Lysak et al. 2016; Mandáková et al. 2020). In *Cucumis* L., genomic blocks combined with FISH maps revealed mechanisms of genome reshuffling, centromere repositioning and decreasing dysploidy in cucumber (*C. sativus* L.) based on ancestral karyotype as reference (Yang et al. 2014; Zhao et al. 2021). Similar ancestral karyotype reconstructions were performed in *Apium* L. (Song et al. 2021), *Camelina* Crantz (Zhang et al. 2020b) and *Cocos* L. (Wang et al. 2021b). However, this requires genome assemblies in chromosomal level and/or chromosome painting probes, which are still not a reality for most group of plants.

Leguminosae L. is one of the largest families of flowering plants. The Papilionoideae subfamily has many economically important members with assembled genome, as the polyploids peanut (2*n* = 4*x* = 40, *Arachis hypogaea* L.), soybean (2*n* = 4*x* = 40, *Glycine max* L.) and white lupin (2*n* = 6*x* = 50, *Lupinus albus* L.), and the diploids common bean (2*n* = 2*x* = 22, *Phaseolus vulgaris* L.), cowpea [2*n* = 2*x* = 22, *Vigna unguiculata* (L.) Walp.] and barrel medic (2*n* = 2*x* = 16, *Medicago truncatula* Gaertn) (Schmutz et al. 2010, 2014; Pecrix et al. 2018; Zhuang et al. 2019; Lonardi et al. 2019; Hufnagel et al. 2020). Comparative genomics revealed a legume-common tetraploidization (LCT) around 60 million years ago (Mya) and independent polyploidization events in *Arachis* (~0.4 Mya), *Glycine* (~ 12 Mya) and *Lupinus* clades (~ 20 Mya) (Schmutz et al. 2010; Wang et al. 2017; Kreplak et al. 2019; Zhuang et al. 2019; Hufnagel et al. 2020). Furthermore, high levels of synteny with large conserved genomic blocks were identified among these genomes (Wang et al. 2017), enabling the reconstruction of ancestral legume karyotypes (ALK), ranging from *n* = 9 (Ren et al. 2019) to *n* = 25 (Kreplak et al. 2019), although *n* = 16 seems more consistent for the ALK (Zhuang et al. 2019; Hufnagel et al. 2020). Most of these analyses used orthologous genes from *P. vulgaris* to detect shared genomic blocks (GBs) among these legumes, indicating preservation of *P. vulgaris* genome (with at least five highly conserved chromosomes) from the legumes ancestor.

Within *Phaseolus* L. genus, BAC-FISH based on *P. vulgaris* probes demonstrated conserved synteny among three species (*P. vulgaris, P. lunatus* L. and *P. microcarpus* Mart.) belonging to different clades (Bonifácio et al. 2012; Fonsêca and Pedrosa-Harand 2013). On the other hand, similar comparative cytogenetic mapping of *P. leptostachyus* Benth. and *P. macvaughii* Delgado (Leptostachyus clade) showed extensive genome reshuffling associated with descending dysploidy involving a nested chromosome fusion between chromosomes 10 and 11 (Fonsêca et al. 2016; Ferraz et al. 2020). Furthermore, comparative cytogenetics and sequence alignment between *Vigna* Savi species and *P. vulgaris* revealed a high degree of macrosynteny between genera, with five chromosomes involved in synteny breaks (Vasconcelos et al.2015; Lonardi et al. 2019; Oliveira et al. 2020; do Vale Martins et al. 2021). A detailed analysis of chromosomes 2 and 3 of *V. angularis, V. unguiculata* and *P. vulgaris* based on sequence alignment and oligo-FISH painting integrative approaches identified additional macro- and micro inversions, translocations, and intergeneric centromere repositioning among analysed species (do Vale Martins et al. 2021). Additionally, centromere repositioning was detected for chromosomes 5, 7, and 9 of *V. unguiculata* and *P. vulgaris* by oligo-FISH barcode combined with genome sequence data (de Oliveira Bustamante et al. 2021). In addition, the authors identified the participation of chromosome 5 in the translocation complex involving chromosomes 1, 5 and 8, a paracentric inversion on chromosome 10, and a pericentric inversion on chromosome 4. The direction and the divergence time of these rearrangements were, however, not estimated.

Although many ALKs have been reconstructed, there is no applicable GB system, which could provide a phylogenetic direction and details of the chromosome rearrangements during legume evolution. Thus, to understand the dynamics of genome reshuffling among the diploid species of Phaseoleae tribe (Leguminosae; Papilionoideae), we constructed a GB system based on comparative cytogenomic data. We compared *P. vulgaris* and *V. unguiculata* genomes to define the GB system using *P. vulgaris* as reference and applied the GBs to four species with assembled genome: *Phaseolus lunatus* L. (2*n* = 2*x* = 22), from the Phaseolinae subtribe; *Spatholobus suberectus* Dunn (2*n* = 2*x* = 18) and *Amphicarpaea edgeworthii* Benth. (2*n* = 2*x* = 22), both belonging to the Glycininae subtribe; and *Medicago truncatula* Gaertn (2*n* = 2*x* = 16, tribe Trifolieae) as an outgroup. Moreover, we performed oligo-FISH using specific probes for *P. vulgaris* chromosomes 2 and 3 to visualize the orthologous chromosomes in two non-sequenced Phaseolinae species with intermediate phylogenetic positions, namely *Macroptilium atropurpureum* DC. Urb. (2*n* = 2*x* = 22) and *Lablab purpureus* L. (2*n* = 2*x* = 22). We hypothesized the Ancestral Phaseoleae Karyotype (APK) and inferred the main chromosomal rearrangements related to the evolution and diversification of these legumes. Our results indicated extensive genome reshuffling in particular lineages and centromere repositioning across the Phaseolinae and Glycininae subtribes at diploid level, with centromere repositioning for all the 11 chromosomes of *P. vulgaris/V. unguiculata*, despite their conserved karyotypes. Our GB system and the proposed APK are promising tools for future comparative genomics analyses when further legumes genome assemblies become available.

## Materials and Methods

### Genomic data sets

We selected and analysed the reference genomes of *P. vulgaris* ‘G19833’ (GenBank ID: 8715468, Schmutz et al. 2014), *P. lunatus* ‘G27455’ (GenBank ID: 20288068, Garcia et al. 2021), *V. unguiculata* ‘IT97K-499-35’ (GenBank ID: 8372728, Lonardi et al. 2019), *S. suberectus* ‘SS-2018’ (GenBank ID: 8715468, Qin et al. 2019) and *A. edgeworthii* ‘Qianfo Mountain’ (GenBank ID: 22470258, Liu et al. 2020). Moreover, we used the *M. truncatula* ‘Jemalong A17’ (GenBank ID: 7445598, Pecrix et al. 2018) genome as an outgroup.

### Genomic blocks definition and APK reconstruction

To identify the syntenic blocks between *P. vulgaris* and *V. unguiculata* genomes, we used the CoGe SynMap platform (https://genomevolution.org/coge/SynMap.pl) (Lyons et al. 2008). We followed the steps and parameters described by Walden et al. (2020): 1) the orthologs were identified by using BlastZ tool; 2) synteny analysis was performed by using DAGChainer, with 25 genes as the maximum distance between two matches (-D), and 20 genes as the minimum number of aligned pairs (-A); 3) Quota Align Merge was used to merge the syntenic blocks, with 50 genes as the maximum distance between them; and 4) the ortholog and paralog blocks were differentiated based on the synonymous substitution rate (Ks) by CodeML (where 2 was the maximum value of log10) and represented by different colours in the dot plot.

The syntenic blocks were defined using the ‘Final syntenic gene-set output with GEvo link’ (Supplementary Table 1), with start and end of the blocks represented by the gene IDs of each species. Each generated genomic block (GB) contained at least 20 genes. The main criterion for the definition of a GB was the presence of translocations break points in its borders.

Based on the gene orthology between *P. vulgaris* and *V. unguiculata* and the defined GBs, we compared the *P. vulgaris* genome with *P. lunatus, S. suberectus, A. edgeworthii*, and *M. truncatula* genomes following the parameters described above. The comparisons generated sub-genomic blocks (sub-GBs) based on the following criteria: 1) breaks of collinearity inside the GBs (inversions); and 2) breaks of collinearity or synteny involving association to other GB (inter- and intrachromosomal translocations). Each species had specific sub-GBs indicating lineage-specific chromosomal rearrangements.

To infer the APK, we analysed all the dotplot patterns and selected the most frequent GB associations, especially those shared with the outgroup (*M. truncatula*), considering the phylogenetic relationships among species, as described by Li et al. (2013), and checking the same or similar breaks points. Altogether, we selected the most conserved GBs and placed them into 11 chromosomes, following the pseudomolecules of *P. vulgaris* genome. Once we defined the APK, we named the GBs in all species using APK as reference.

The centromeric regions were defined based on the centromeric data available in the genome assemblies (Schmutz et al. 2014; Pecrix et al. 2018; Lonardi et al. 2019; Liu et al. 2020; Garcia et al. 2021). As the centromere positions for *S. suberecuts* were not indicated in the genome assembly, we hypothesized the centromere region according to the peaks of TE accumulation along the chromosomes (Qin et al. 2019). Finally, we standardize the centromere regions using the mid-point of each centromeric region, considering that each centromere represents ~ 2 Mb.

### Plant material and chromosome preparation

For the cytogenetics analyses, we used *P. vulgaris* ‘BAT 93’ (Embrapa Recursos Genéticos e Biotecnologia - Cenargen, Brasília, Distrito Federal, Brazil), *V. unguiculata* ‘BR14 Mulato’ (Embrapa Meio-Norte, Teresina, Piauí, Brazil), *M. atropurpureum* (International Center for Tropical Agriculture, CIAT 4413) and *L. purpureus* (UFP87699). Root tips from germinated seeds were collected and pre-treated with 2 mM 8-hydroxyquinoline for 5 h at 18° C, fixed in methanol or ethanol: acetic acid (3:1 v/v) for 2-24 h at room temperature and stored at −20 °C until use. For chromosome preparation, the roots were washed twice with distilled water, digested with an enzymatic solution containing 2% pectolyase (Sigma-Aldrich), 4% cellulase (Onozuka or Sigma-Aldrich) and 20% pectinase (Sigma-Aldrich) for 1-2 h at 37°C (humid chamber). Slides were prepared following the air dry protocol (Carvalho and Saraiva 1993), with minor modifications.

### Oligo-FISH, image acquisition and data processing

The design, synthesis and labelling of the *Pv*2 and *Pv*3 oligo probes were described by do Vale Martins et al. (2021). Oligo-FISH was carried out according to Han et al. (2015), with minor changes. The hybridization mixture consisted of 50% formamide, 10% dextran sulphate, 2× saline sodium citrate (SSC), 350 ng of the biotin-labelled probe (*Pv*2, green) and 300 ng of the digoxigenin-labelled probe (*Pv*3, red), in a total volume of 10 μL per slide. The hybridization mix was applied to the slides for 5 min at 75°C and hybridized for 2-3 days at 37°C. *Pv*2 and *Pv*3 oligo probes were detected with anti-biotin fluorescein (Vector Laboratories) and anti-digoxigenin rhodamine (Roche), respectively, both diluted in 1× TNB (1M Tris HCl pH 7.5, 3 M NaCl-blocking reagent, Sigma-Aldrich) with posterior incubation for 1 h at 37°C. Chromosomes were counterstained with 2 μg/mL DAPI in Vectashield antifade solution (Vector Laboratories). Oligo-FISH images were captured with a Hamamatsu CCD camera attached to an Olympus BX51 epifluorescence microscope or Leica DM5500B fluorescence microscope. The images were adjusted and optimized for brightness and contrast using Adobe Photoshop CC (2019). *Phaseolus vulgaris* and *V. unguiculata* idiograms were assembled based on Vasconcelos et al. (2015).

### Accession Numbers

*Phaseolus vulgaris* ‘G19833’ (GenBank ID: 8715468, Schmutz et al. 2014)

*Phaseolus lunatus* ‘G27455’ (GenBank ID: 20288068, Garcia et al. 2021),

*Vigna unguiculata* ‘IT97K-499-35’ (GenBank ID: 8372728, Lonardi et al. 2019),

*Spatholobus suberectus* ‘SS-2018’ (GenBank ID: 8715468, Qin et al. 2019)

*Amphicarpaea edgeworthii* ‘Qianfo Mountain’ (GenBank ID: 22470258, Liu et al. 2020)

*Medicago truncatula* ‘Jemalong A17’ (GenBank ID: 7445598, Pecrix et al. 2018)

## RESULTS

### Genomic blocks and the inferred Ancestral Phaseoleae Karyotype

To infer the ancestral karyotype of the Phaseoleae tribe, we aligned *P. vulgaris* (*Pv*) and *V. unguiculata* (*Vu*) genomes based on the collinear arrangement of orthologous genes in dot plots (Supplementary Table 1) following Walden et al. (2020). Then, we compared the *Pv* genome to *P. lunatus* (*Pl*), *S. suberectus* (*Ss*), *A. edgeworthii* (*Ae*) and *M. truncatula* (*Mt*) genomes, in order to detect the synteny blocks (Supplementary Table 2-5). We selected the most frequent GB associations, particularly those shared with *M. truncatula* (outgroup), considering the phylogenetic relationships among species, as described by Li et al. (2013), and checking if they shared the same or similar break points by the dot plot patterns (Supplementary Figures 1-5). We proposed the APK with *n* = 11 (most common chromosome number within the tribe; Rice et al. 2015) and 19 GBs (A-S), ordered according to the pseudomolecules of the *Pv* genome (Figure 1A; Supplementary Table 6). Due to inversions within a block or transpositions involving part of the blocks, some blocks were divided into sub-blocks (Supplementary Table 7-12). Different numbers of sub-blocks were defined for each species, due to independent rearrangements among them. The centromere positions in APK chromosomes were hypothesized based on the frequency of associations between GBs and centromeres in the analysed species, although some of them might not represent the ancestral state.

**Figure 1.**
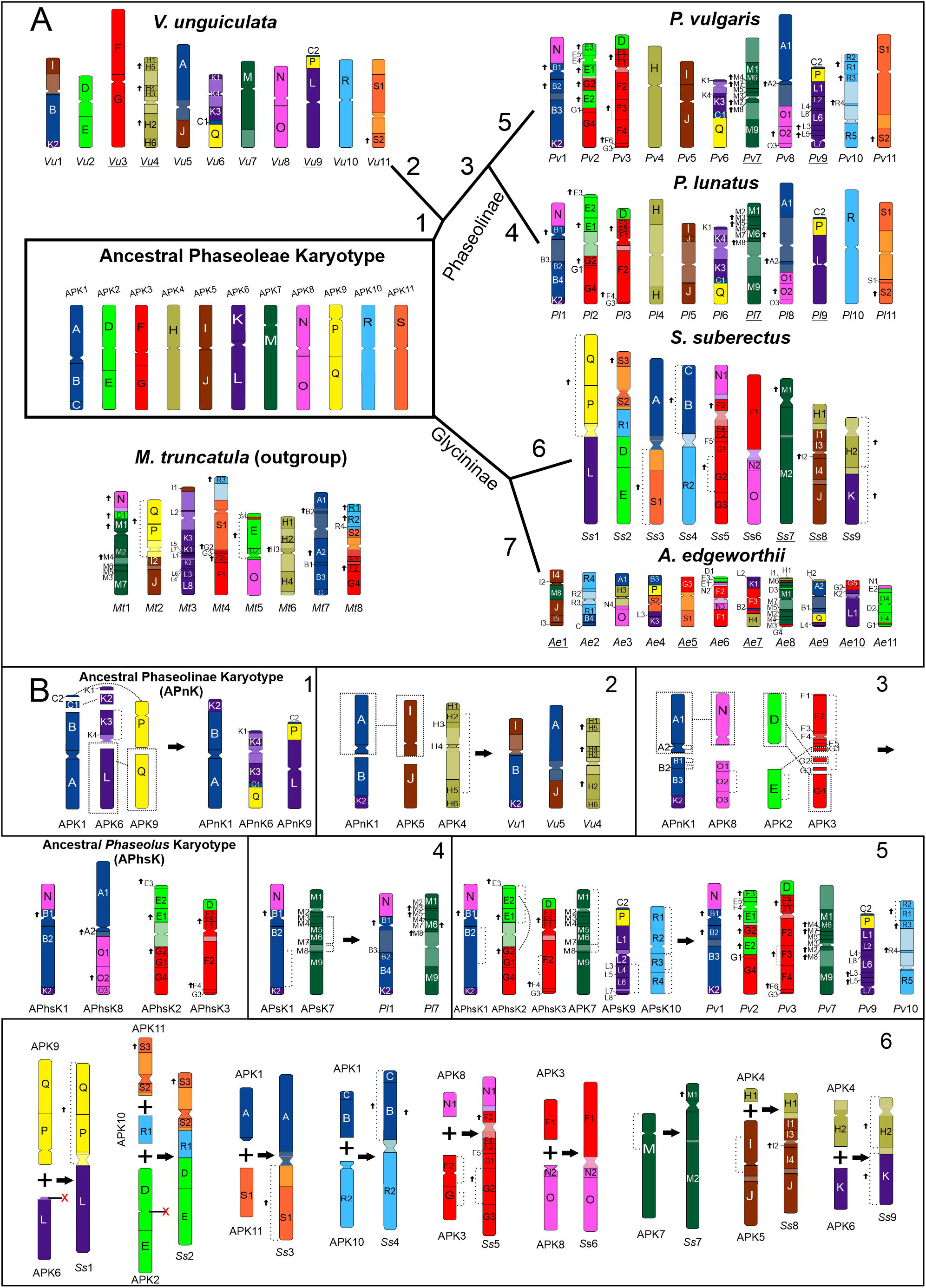
Genomic block system and the hypothetical Ancestral Phaseoleae Karyotype (APK). Arrows in **A** indicate change in block orientation. Some entire chromosome orientations were inverted (underlined chromosomes) to better demonstrate the orthology between APK and other karyotypes. **A**) Species are grouped according to the phylogenetic relationships proposed by Li *et al*. (2013). **1** to **5**) Phaseolinae subtribe species: *Vigna unguiculata, Phaseolus lunatus* and *P. vulgaris*; **6** and **7**) Glycininae subtribe species: *Spatholobus suberectus* and *Amphicarpaea edgeworthii*. **B**) Main chromosomal rearrangements: **1**) Translocations and pericentric inversion resulting in the Ancestral Phaseolinae Karyotype (APnK); **2**) Reciprocal translocation and pericentric inversions resulting in *V. unguiculata* (*Vu*) chromosomes 1, 4 and 5; **3**) Reciprocal translocation and inversions resulting in the Ancestral *Phaseolus* Karyotype (APhsK); **4**) Inversions and intrachromosomal translocations resulting in *P. lunatus* (*Pl*) chromosomes 1 and 7; **5**) Inversions and intrachromosomal translocation resulting in *P. vulgaris* (*Pv*) chromosomes 1, 2, 7, 9 and 10; **6**) Descending dysploidy and genome rearrangements for *S. suberectus* (*Ss*) karyotype.

### Chromosomal rearrangements and centromere repositioning in Phaseolinae subtribe in relation to the APK

Eight APK chromosomes displayed full synteny with at least one chromosome of the three analysed Phaseolinae subtribe species: APK2 (*Vu*2), APK3 (*Vu*3), APK4 (*Pv*4, *Pl*4, *Vu*4), APK5 (*Pv*5, *Pl*5), APK7 (*Vu*7, *Pv*7, *Pl*7), APK8 (*Vu*8), APK10 (*Pv*10, *Pl*10, *Vu*10) and APK11 (*Pv*11, *Pl*11, *Vu*11). We propose that the main chromosomal rearrangements common to all Phaseolinae species (Figure 1.B1) involved a complex translocation among APK1, 6 and 9, resulting in the ancestral Phaseolinae karyotype (APnK) chromosomes 1, 6 and 9 (Figure 1.B1). We confirmed exclusive rearrangements for *V. unguiculata* and *Phaseolus* species (Figure 1.B2 and 1.B3, respectively). In *V. unguiculata* (*Vu*), we observed a reciprocal translocation between APnK1 and APK5, resulting in chromosomes *Vu*1 (I+B+K2) and *Vu*5 (A+J), in addition to a large pericentric inversion comprising most of APK4 (H), corresponding to *Vu*4 (Figure 1.B2). Furthermore, in the ancestral *Phaseolus* karyotype (APhsK), two reciprocal translocations occurred between APnK1 and APK8, resulting in APhsK1 (N+B+K2) and 8 (A+O); and between APK2 and 3, generating APhsK2 (E+G) and 3 (D+F), followed by inversions and intrachromosomal translocations on APhsK2 (E and G) and 3 (F) (Figure 1.B3). We also detected previously described inversions between *P. vulgaris* and *P. lunatus* (Bonifácio et al. 2012; Garcia et al. 2021) in Chr1 (in B); Chr2 (E and G); Chr3 (within F2); Chr7 (M); Chr9 (in L), and Chr10 (in R). The complex multiple inversions in chromosomes 2 and 7, also involving intrachromosomal translocations, occurred independently in *P. vulgaris* and *P. lunatus*, while the intrachromosomal translocations on chromosome 1 and inversions on chromosomes 1, 3, 9 and 10 occurred in *P. lunatus* (*Pl*1) and *P. vulgaris* (*Pv*1, 3, 9 and 10) lineages, respectively (Figure 1.B4 and 1.B5).

Our analyses indicated centromere repositioning in at least one of the analysed Phaseolinae species based on centromere position relative to the GBs (Figure 2). The more evident repositioning events were those in which the centromere position changed the GB association (APK2: *Pl*/*Pv*2; APK3: *Vu*3; APK5: *Pl*/*Pv*5; APK6: *Vu*/*Pl*/*Pv*6; APK8: *Pl*/*Pv*1; APK9: *Vu*/*Pl*/*Pv*9; Figure 2). Some centromeres were preserved on the same GBs; however, sub-GBs showed that the positions were apparently changed (APK7: *Pl*/*Pv*7; APK10: *Pv*10), which could indicate further centromere repositioning events (Figure 2). To further investigate these additional putative centromere repositioning events, we compared the syntenic genes blocks flanking the centromere regions in *Phaseolus* and *Vigna* (gene borders in Supplementary Table 7 - 9). The results demonstrate that all gene borders of centromeres had significant differences, indicating centromere repositioning in all centromeres between *P. vulgaris* and *V. unguiculata*. The majority of these repositioning events probably occurred in *P. vulgaris* lineage, since the major divergence of GBs and of genes borders among species was detect in *P. vulgaris* (Figure 2; Supplementary Table 7 – 12). Overall, we hypothesize that the major events of centromere repositioning were derived from Evolutionary New Centromere (ENC) events because they are present at different GBs or sub-GBs without associated chromosomal rearrangements (Figure 2).

**Figure 2.**
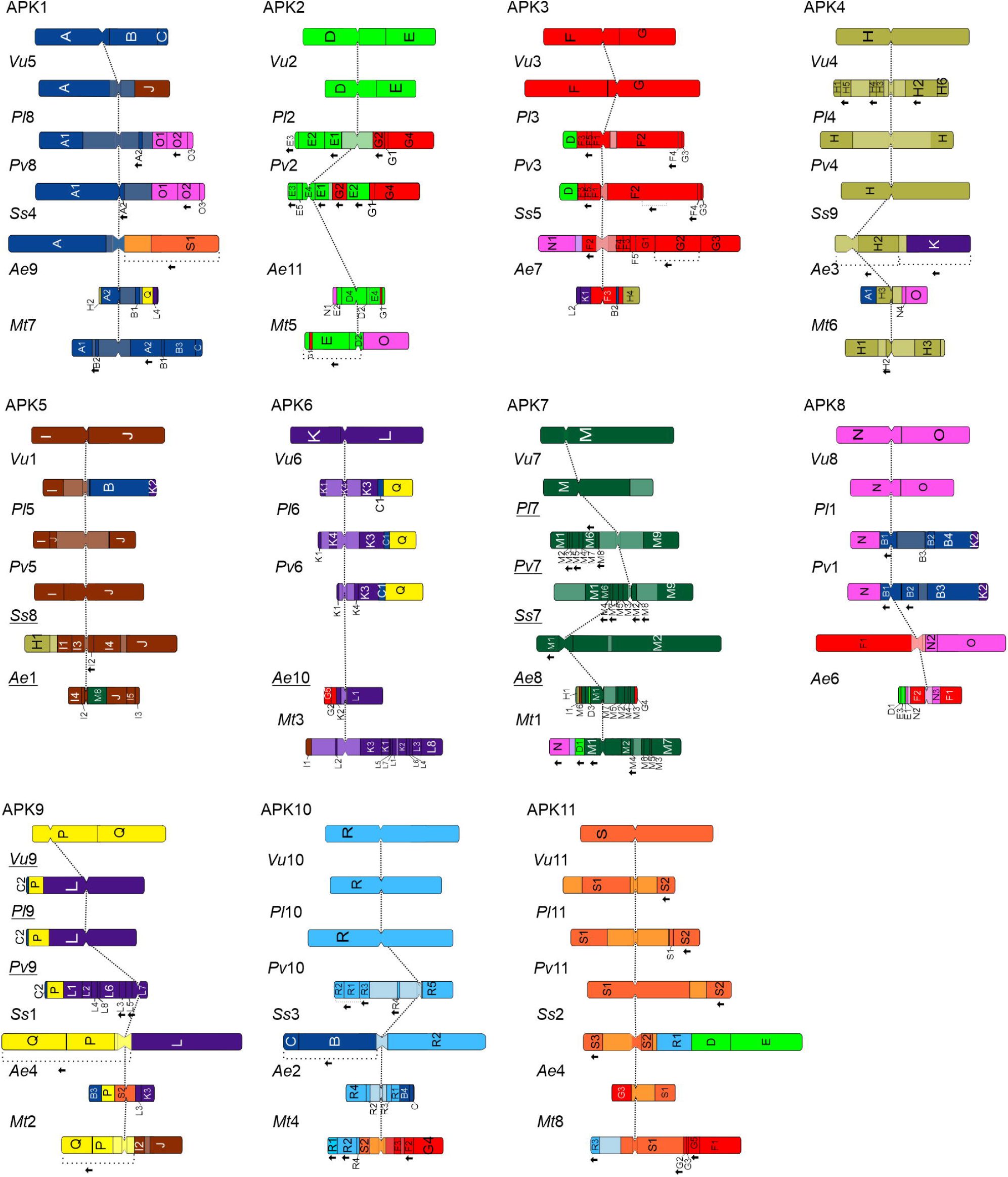
Centromere repositioning in Phaseoleae tribe and *M. truncatula* based on the established Ancestral Phaseoleae Karyotype (APK).

### The main rearrangements among Phaseolinae, Glycininae and *M. truncatula* inferred from comparison with APK

All the generated GBs were conserved in *S. suberectus, A. edgeworthii* (Glycininae) and *M. truncatula* (Figure 1A). Two APK associations were shared between *S. suberectus* and *M. truncatula:* B+C (APK1: *Ss*4, *Mt*7) and Q+P (APK9: *Ss*1, *Mt*2). However, these associations were not evidenced within the Phaseolinae species. The GBs M (APK7: *Pv*7, *Pl*7, *Vu*7, *Ss*7, *Mt*1) and O (APK8: *Pv*8, *Pl*8, *Vu*8, *Ss*6, *Ae*3, *Mt*5) were highly syntenic between the subtribes, with an exclusive inversion in O2 in *Phaseolus* (Figure 1A). The GB M was also maintained in a single chromosome among species, except for *A. edgeworthii* (M8: *Ae*1 and M1-M7: *Ae*8; Figure 1A).

Comparison between Glycininae and Phaseolinae subtribes showed that almost all chromosomes were involved in breaks of synteny and/or collinearity, which led to a higher number of sub-GBs, especially in *A. edgeworthii*. Although this species maintained the ancestral chromosome number *n* = 11, several rearrangements lead to complex GB associations (Figure 1.A7). Nevertheless, we were able to find the APK associations that helped to unveil the main translocations in this genome (Supplementary Figure 6). On the other hand, despite the descending dysploidy to *n* = 9, the *S. suberectus* genome showed fewer rearrangements when compared to the APK than the Phaseolinae species (Figure 1.A6). All the 19 GBs were detected in *M. truncatula* genome (which diverged from *P. vulgaris* ~50 Mya), with *Mt*1 (M) and *Mt*6 (H) highly syntenic to Phaseolinae chromosomes 7 and 4, respectively (Figure 1A).

Based on the APK, we proposed that the main chromosomal rearrangements that lead to the descending dysploidy in *S. suberectus* (*n* = 11 to *n* = 9) were due to chromosome fusions. These rearrangements involved seven APK chromosomes (APK2, APK4, APK5, APK6, APK9, APK10 and APK11), resulting in four *S. suberectus* chromosomes (*Ss*1, *Ss*2, *Ss*8 and *Ss*9) (Figure 1.B6). APK 4, 5, 6 and 9 were involved in a complex translocation, originating *Ss*1, *Ss*8 and *Ss*9. The whole APK2 and part of APK10 and APK11 were combined by a translocation with terminal breakpoints, resulting in *Ss*2, followed by centromere loss in the D block. Additional reciprocal translocations occurred between APK1 with APK10 and APK11 generating *Ss*3 and *Ss*4, and between APK3 and APK8, resulting in *Ss*5 and *Ss*6. APK7 was conserved in *S. suberectus*.

### Reconstructing the chromosomal rearrangements in APK based on the Ancestral Legume Karyotype

Using data from ALK and the rearrangements in *P. vulgaris* detected by Hufnagel et al. (2020), we identified the corresponding blocks from three ancestral karyotypes of legumes in our proposed APK (Supplementary Table 3). We hypothesize that the formation of APK involved five chromosomal fusions between 10 chromosomes of ALK, which reduced the chromosome number from *n* = 16 to *n* = 11 (Figure 3). Considering ALK’s from Hufnagel et al. (2020), Kreplak et al. (2019) and Zhuang et al. (2019), we observed that at least six APK chromosomes were well preserved during the diversification of legumes (APK4, 5, 7, 9, 10 and 11), what explains the preservation of these chromosomes in many analysed species (Figure 1; Supplementary Table 3). The APK patterns for *P. vulgaris* and *M. truncatula* are similar to the ALK pattern (Hufnagel et al. 2020), corroborating our APK reconstruction. Moreover, the APK pattern for *M. truncatula* confirms the translocation between *Mt*4-*Mt*8 (Kamphuis et al. 2007), which involved APK3, 10 and 11.

**Figure 3.**
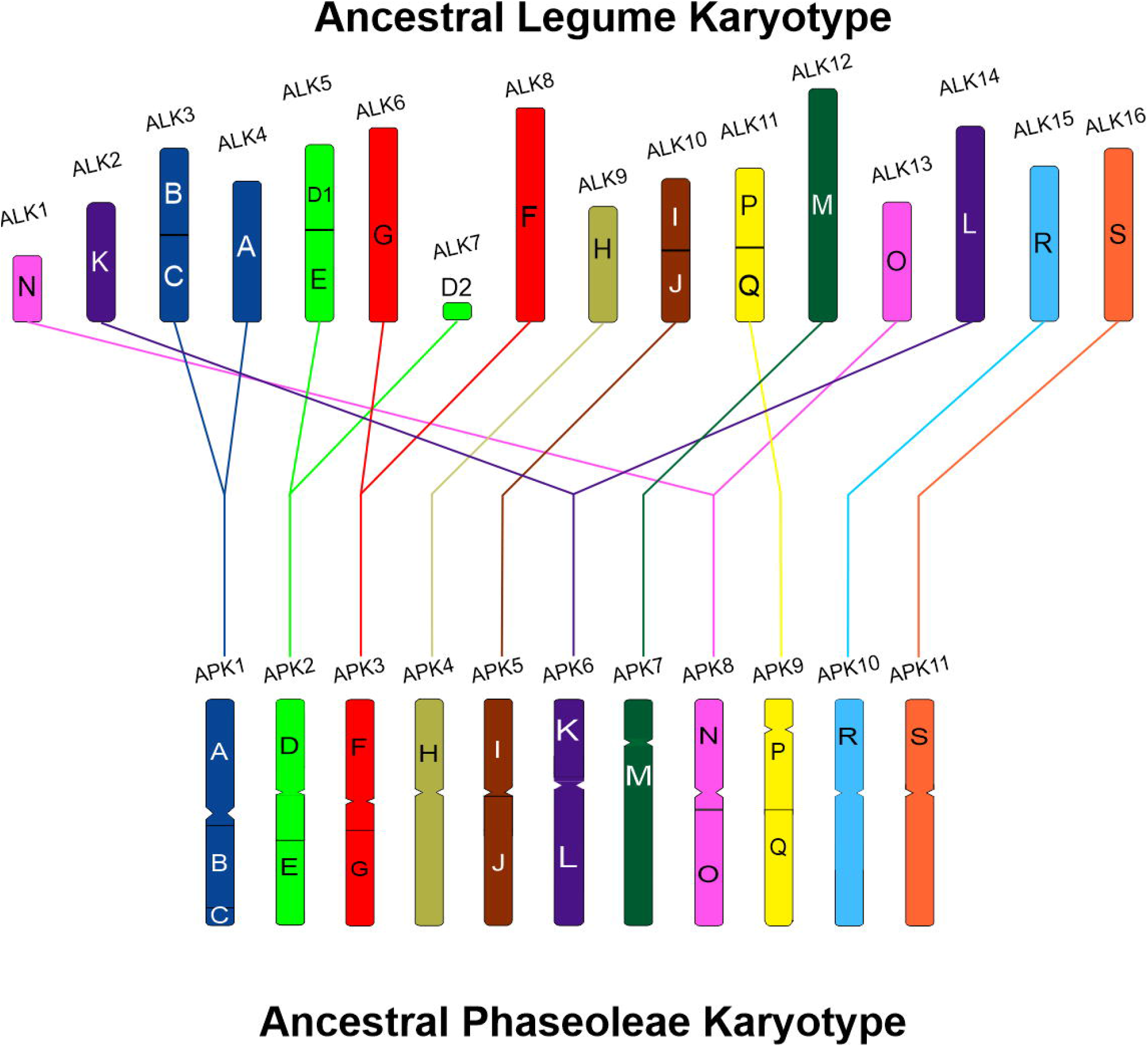
Ancient Legume Karyotype (ALK, *n* = 16) by Hufnagel et al. (2020) compared to the APK (*n* = 11). Ten ALKs were involved in five end-to-end chromosome fusions, reducing from *n* = 16 to *n* = 11.

### Oligopainting of chromosomes 2 and 3 in the two non-sequenced species *M. atropurpureum* and *L. purpureus*

To further investigate the chromosome evolution within the Phaseolinae subtribe, we expanded our comparative cytogenomic analysis in two species with no assembled genome, namely *M. atropurpureum* (*Ma*) and *L. purpureus* (*Lp*). We hybridized two oligopainting probes from *P. vulgaris* chromosomes (Figure 4a), *Pv*2 (green) and *Pv*3 (red) to *M. atropurpureum* and *L. purpureus* metaphase cells. The oligo-FISH painting did not reflect the patterns expected for APK because the probes were generated from *Pv* chromosomes, which experienced rearrangements between chromosomes 2 and 3 (Figure 4). *Macroptillium atropurpureum* (Figure 4c) and *L. purpureus* (Figure 4d) showed similar oligo-FISH signals to *V. unguiculata* (Figure 4b). *Macroptillium atropurpureum* ortholog chromosome 2 (*Ma*2) presented the short arm in red (*Pv*3) and almost the entire long arm in green (*Pv*2), except for an interstitial pericentromeric red region on its long chromosome arm. The *Ma*3 presented the short arm and about half of the long arm in red (*Pv*3), with the distal region painted in green (*Pv*2) (Figure 4c). In the *L. purpureus* (Figure 4d) chromosomes, we evidenced a more complex oligopainting pattern: the short arm of chromosome 2 (*Lp*2) showed small terminal and pericentromeric green (*Pv*2) signals intermingled with a large red (*Pv*3) signal, while the opposite arm, similar to *Vu*2 long arm, was painted in green (*Pv*2), with a pericentromeric region in red (*Pv*3). The *Lp*3 short arm, similar to the short arm of *Vu*3, was painted in green (*Pv*2), with an intermingled small red signal (*Pv*3). The long arm of *Lp*3 was painted in red (*Pv*3), with an intermingled green signal (*Pv*2) in the proximal region of its arm (Figure 4d).

**Figure 4.**
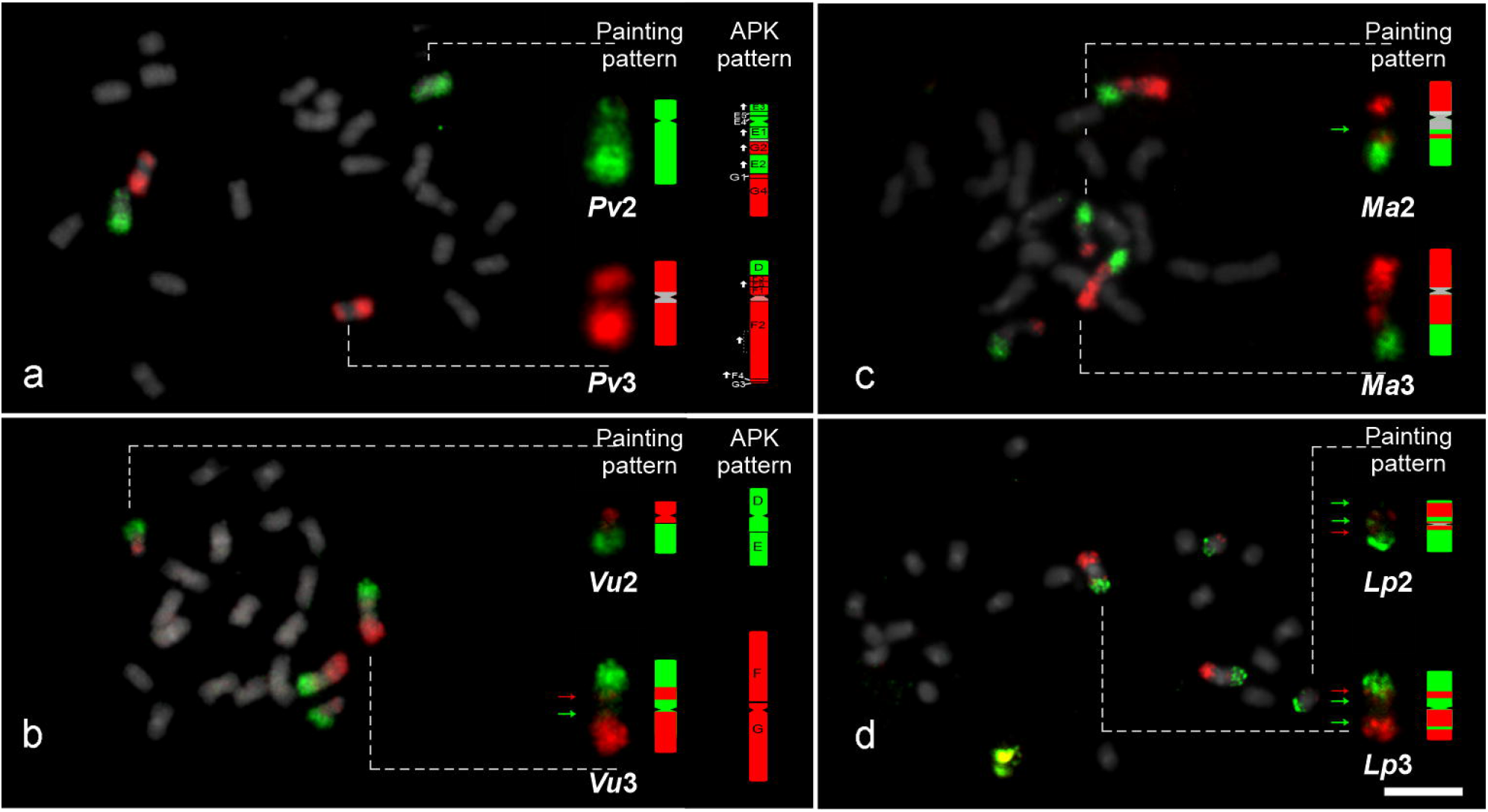
Oligo-FISH using *Phaseolus vulgaris* chromosomes-specific 2 (green) and 3 (red) probes hybridized to other Phaseolinae species. (**a**) *P. vulgaris*, (**b**) *Vigna unguiculata*, (**c**) *Macroptilium atropurpureum* and (**d**) *Lablab purpureus*. Green and red arrows indicate small differences in the painting patterns, corresponding to specific regions of *Pv*2 (green) or *Pv*3 (red) probes, respectively. For each species, orthologous chromosomes of *P. vulgaris* chromosomes 2 and 3 are detailed in the insets (right) of each metaphase cell and represented on the right of each inset. Differences between the APK and oligo-FISH painting signals of chromosomes 2 and 3 are indicated for *P. vulgaris* and *V. unguiculata*. Vertical bar = 5 μm.

Our data support the exclusivity of the translocation event between APK2 and APK3 for the genus *Phaseolus*, since *L. purpureus* and *M. atropurpureum* chromosomes 2 and 3 resemble *V. unguiculata* ortholog chromosomes and are closer to the APK. However, gaps in the pericentromeric regions of *M. atropurpureum* and *L. purpureus* chromosomes 2 and 3 (Figure 4c, 4d), different chromosome arm sizes in *Ma*3 (Figure 4c), and intermingled oligo-FISH signals in *L. purpureus* (represented by green and red arrows in Figure 4d), may indicate independent rearrangements and small breaks of collinearity that can be related to inversions.

## Discussion

Here we established a GB/sub-GB system for comparative chromosome analyses of Phaseoleae legumes. Our system identified several chromosomal rearrangements and frequent centromere repositioning, especially in the *Phaseolus* lineage. The GB system was also applicable for *Medicago* (distantly related tribe), confirming the high synteny inside the legume family and suggesting its use to unveil chromosome evolution in a wide range of legumes with sequenced genomes. The sub-GBs revealed further rearrangements inside the GBs as independent events during evolution. In Brassicaceae, the identification of independent rearrangements inside the GBs was essential for understanding the phylogenetic relationships in different taxa, such as *Aethionema arabicum* (L.) Andrz. ex DC. (Walden et al. 2020), *Arabis alpina* L. (Willing et al. 2015) and *Brassica oleracea* L. (Parkin et al. 2014). The proposed GB system might be useful in future phylogenetic analyses within the Leguminosae family.

Recent comparative genomics were able to reconstruct the ALKs, with proposed chromosome numbers ranging from *n* = 9 to *n* = 25 (Zhuang et al. 2019; Kreplak et al. 2019; Ren et al. 2019; Hufnagel et al. 2020). Nevertheless, comparisons among grape (*Vitis vinifera* L.) and legumes revealed independent chromosome fusions before and after the LCT event, resulting in *n* = 16 as the putative ancestral chromosome number for legumes (Zhuang et al. 2019). The ALKs unveiled detailed chromosome rearrangement events that shaped the chromosome number in specific lineages, mainly marked by polyploidization and chromosome fusions and fissions (Zhuang et al. 2019; Kreplak et al. 2019; Hufnagel et al. 2020). Additional analyses, as centromere repositioning and inversions, were not discussed, and *V. unguiculata, Phaseolus lunatus*, *Spatholobus suberectus* and *Amphicarpaea edgeworthii* were not investigated (Zhuang et al. 2019; Kreplak et al. 2019; Hufnagel et al. 2020).

We reconstructed an ancestral Phaseoleae karyotype (APK), with *n* = 11 chromosomes and 19 GBs. Despite the chromosome number variation inside the tribe (2*n* = 18 to 2*n* = 84; Rice et al. 2015), genomic, cytogenetic and phylogenetic evidences suggest the ancestral chromosome number as *n* = 11 (Li et al. 2013; Rice et al. 2015; Wang et al. 2017). Comparing the ALK (Hufnagel et al. 2020) with our APK, we were able to point the chromosome number reduction of *n* = 16 to *n* = 11, which involved five chromosome end-to-end fusions. Reconstruction of ancestral karyotypes has been essential for comparative genomic analyses in other groups, as we see in grasses, cucurbits, crucifers and other flowering plants, contributing to comprehend the chromosome number variation, genome reshuffling and recombination hotspots (Murat et al. 2010, 2017; Lysak et al. 2016; Xie et al. 2019). More recently, an ancestral karyotype of *Cucumis* was inferred by comparative oligo-painting (COP) in different species of African and Asian clades, indicating constant genome reshuffling caused by large-scale inversions, centromere repositioning, and other additional rearrangements (Zhao et al. 2021). Similarly, our analyses showed highly conserved macrosynteny in the Phaseoleae tribe, revealing the particular rearrangements within each clade.

Five APK chromosomes (APK4, APK5, APK7, APK10 and APK11) showed high conservation of synteny within the tribe and with the ALK (Hufnagel et al. 2020), as observed in previous studies (Schmutz et al. 2010; McConnell et al. 2010; Wang et al. 2017; Ho et al. 2017; Lonardi et al. 2019). Overall, APK7 is the most conserved chromosome, with the GB “M” involved only in intrachromosomal rearrangements, except for *A. edgeworthii*, which displayed a higher number of rearrangements. Chromosome 7 showed high conservation of synteny when compared to the non-Phaseoleae species, such as *Mt*1, *Ah*9 (*Arachis hypogaea* L. chromosome 9) and *Lj*5 [*Lotus japonicus* (Regel) K.Larsen chromosome 5] (Bertioli et al. 2009), as well as to soybean *Gm*10 and *Gm*20 chromosomes (Schmutz et al. 2014; Wang et al. 2017). Gene family analyses in *Pisum sativum* L. and *M. truncatula* indicate expression of important genes for the seed storage (Legumin, Vicilin and Convicilin) in the *Ps*6 and *Mt*1, orthologous to APK7 (Kreplak et al. 2019). This could indicate that the high synteny of this chromosome may be important for seed development. Additional analyses are necessary to test this hypothesis.

Few translocations and a large number of inversions were identified within Phaseolinae. Some of these rearrangements were previously identified by BAC-FISH, oligo-FISH and comparative genomics in *P. vulgaris, P. lunatus* and *V. unguiculata* (Bonifácio et al. 2012; Vasconcelos et al. 2015; Lonardi et al.2019; Oliveira et al. 2020; do Vale Martins et al. 2021; Garcia et al. 2021; de Oliveira Bustamante et al.2021). Based on our APK and oligo-FISH approaches, we are now able to propose the direction of these rearrangements in a phylogenetic context. The reciprocal translocation between chromosomes 1 and 8, 2 and 3, and inversions on chromosomes 2 and 3 were exclusive events of *Phaseolus* genus, while the translocation between chromosomes 1 and 5, and the inversions on chromosome 4 were exclusive of *Vigna*. Independent intrachromosomal translocations and inversions in *P. vulgaris* (*Pv*1, 2, 3, 7, 9 and 10) and *P. lunatus* (*Pl*1, 2 and 7) were also detected. Translocations and inversions are important drivers of speciation, meiotic recombination reduction (Rieseberg 2001; Noor et al. 2001; Faria and Navarro 2010; Feulner and De-Kayne 2017) and chromosomal evolution (Martin et al. 2020). Chromosomal inversions in *Drosophila persimilis* and *D. pseudoobscura*, for instance, were associated with reproductive isolation (Fuller et al. 2018).

Several translocations and inversions were detected in the Glycininae subtribe. Although the ancestral chromosome number was conserved in *A. edgeworthii*, its karyotype was the most rearranged among the analysed species. Whole genome analysis indicated large-scale ectopic recombination and reduction of Long Terminal Repeats (LTR) retrotransposons during evolution, compacting the genome but preserving important genes (Liu et al. 2020). The genomic analysis also indicated that remarkable genome reshuffling was not a consequence of polyploidy (Liu et al. 2020). Indeed, none of the GBs were duplicated, but multiple GBs were subdivided due to the extensive rearrangements. On the other hand, the dysploid *S. suberectus* species presented the most conserved karyotype. We proposed two major translocations events combining APK6 and 9 into *Ss*1, and APK2, APK10 and APK11 into *Ss*2, leading to the descending dysploidy in *S. suberectus* (from *n* = 11 to *n* = 9). Centromeres of APK6 and 9 were probably fused in a Robertsonian translocation, while the centromere of APK2 was probably eliminated. Descending dysploidy events were also unveiled by ancestral karyotype analysis in Brassicaceae (Lysak et al. 2006, 2016; Schranz et al. 2006) and in the wild *C. sativus* var*. hardwickii*, from *n* = 12 to *n* = 7 (Yang et al. 2014).

Despite the uncertain centromere position of *S. suberectus* pseudomolecules in the genome assembly, several GB-centromeric associations were conserved among Phaseoleae and in other legumes. Nevertheless, our GBs system revealed that centromere repositioning was common and not associated with other rearrangements or chromosome number reduction, especially in Phaseolinae. We confirmed previously inferred centromere repositioning for five chromosome pairs between *P. vulgaris* and *V. unguiculata*: chromosomes 2, 3, 5, 7 and 9 (do Vale Martins et al. 2021; de Oliveira Bustamante et al.2021). Additionally, we identified centromere repositioning for the remaining six chromosome pairs. Thus, our results indicated repositioning for all centromeres between these two closely related species, separated by ~10 Mya (Li et al. 2013). Apart from one putative event in the ancestral of both genera (in Chr. 9), only one event happened at the *V. unguiculata* lineage, the remaining probably occurring in the lineage that led to *P. vulgaris*.

*Phaseolus vulgaris* and *V. unguiculata* have different centromeric satDNA families and more than one centromeric tandem repeat sequence in each species (Iwata et al. 2013; Iwata-Otsubo et al. 2016). In addition, repetitive sequences showed different amplification dynamics within *Phaseolus* and among *Phaseolus, Vigna* and *Cajanus*, indicating fast evolution of centromere repeats in Phaseoleae (Iwata et al. 2013; Iwata-Otsubo et al. 2016; Ribeiro et al. 2017, 2020). As demonstrated by our GB analyses, repositioned centromeres were not associated to changes in macro-collinearity. These changes may have occurred via invasion of retroelements and tandem repeats, and recruitment of the centromeric protein CENH3 to a new position, forming a functional ENC (Schubert 2018; Talbert and Henikoff 2020), as evidenced in *Solanum* L. (Gong et al. 2012; Zhang et al. 2014), *Oryza* L. (Liao et al. 2018), *Aubrieta* Adans. and *Draba* L. (Mandáková et al. 2020). In maize (*Zea mays* L.), for example, centromere repositioning occurred after DBS in centromeric satellite sequences that led to loss of *CentC*, and resulted in neocentromere formation at linked genes (selected during domestication), which facilitate the emergence of ENC (Schneider et al. 2016; Zhao et al. 2017).

While centromere repositioning in Phaseoleae (especially in Phaseolinae) seems to result from ENC events, chromosome rearrangements, such as inversions and translocations, might have resulted in centromere repositioning (Schubert 2018; Talbert and Henikoff 2020). In *S. suberectus*, two ancestral centromeres seem to be involved in the dysploidy event through a Robertsonian translocation (Figure 1). The association of chromosome rearrangements and centromere repositioning was observed in *Arabis alpina* (Willing et al. 2015), which preserved some associations of centromere-GBs from ACK chromosomes. Altogether, our findings suggest that *de novo* centromere formation is the main mechanism responsible for the frequent centromere repositioning among the GBs observed in Phaseoleae legumes. Further assembly of whole centromeres after long read sequencing will be necessary to confirm the additional role of chromosome rearrangements within GBs and/or repetitive sequence.

Our GBs system enabled us to reconstruct the ancestral karyotype of the Phaseoleae tribe and to determine the direction of the main chromosomal evolutionary events related to diversification and speciation of closely and distantly related legume species. We correlated our APK with proposed ALKs, showing that the chromosome reduction in the tribe was caused by five end-to-end chromosome fusions. At least five legume chromosomes are highly conserved through diversification of the group. Moreover, we observed frequent centromere repositioning in this group, especially in *Phaseolus* genus, despite its karyotype stability. This repositioning likely involved the emergence of ENC. Our findings open a bright perspective for future genomic analysis within Phaseoleae tribe and other legume species. With an increasing number in legume genome sequencing, this genomic tool provides new perspectives for understanding the events that shaped the actual legume genomes, as well as the role of centromere repositioning for plant evolution in a phylogenetic context.

## DECLARATIONS

### Availability of data and material

All data generated or analyzed during this study are included as supplementary materials.

### Code Availability

Not applicable

### Author’s contributions

C.M.: conducted the genome comparison, defined the blocks, performed the oligo-FISH painting experiments in *M. atropurpureum* and *L. purpureus*, constructed the images and wrote the original draft of the manuscript. L.V.M: synthesized the oligo painting probes, performed the oligo-FISH in *P. vulgaris* and *V. unguiculata*, constructed the oligo-FISH image and helped to write the manuscript. F.O.B: provided the resources for this research and discussed the data. A.C.B.V: co-supervised the experiments and contributed with data analyses and discussion. A.P.H: conceptualized and supervised the experiments and provided resources for this research. All authors reviewed the manuscript.

### Ethical approval

Not applicable

### Consent to participate

Not applicable

### Consent for publication

Not applicable

### Key message

We developed a useful genomic block system and proposed the Ancestral Phaseoleae Karyotype (APK) based on available genome assemblies of legume crops. These tools enabled the reconstruction of the main chromosomal rearrangements responsible for the genome reshuffling among the diploid investigated taxa. The analyses revealed centromere repositioning for all chromosomes within the tribe, even when chromosome numbers were conserved.

## Acknowledgments

We thank Embrapa Meio-Norte (Teresina, Piauí, Brazil), Embrapa Cenargen (Brasília, Distrito Federal, Brazil), CIAT (International Center for Tropical Agriculture) and Prof. Marcelo Guerra (UFPE) for providing the *V. unguiculata, P. vulgaris, M. atropurpureum* and *L. purpureus* seeds, respectively. We thank Ingo Schubert (IPK) and André Marques (MPIPZ) for critical early review of the manuscript. We also thank FACEPE (Fundação de Amparo à Ciência e Tecnologia do Estado de Pernambuco) and CNPq (Conselho Nacional de Desenvolvimento Científico e Tecnológico) for financial support.

## SUPPLEMENTARY INFORMATION CAPTIONS

**Supplementary Figure 1.** Dot plot of genome comparison between *P. vulgaris* and *V. unguiculata* with the indication of each GB coloured based on the APK. Corresponding chromosomes are distributed according to dot plot order.

**Supplementary Figure 2.** Dot plot of genome comparison between *P. vulgaris* and *P. lunatus* with the indication of each GB coloured based on the APK. Corresponding chromosomes are distributed according to dot plot order.

**Supplementary Figure 3.** Dot plot of genome comparison between *P. vulgaris* and *S. suberectus* with the indication of each GB coloured based on the APK. Corresponding chromosomes are distributed according to dot plot order.

**Supplementary Figure 4.** Dot plot of genome comparison between *P. vulgaris* and *A. edgeworthii* with the indication of each GB coloured based on the APK. Corresponding chromosomes are distributed according to dot plot order.

**Supplementary Figure 5.** Dot plot of genome comparison between *P. vulgaris* and *M. truncatula* with the indication of each GB coloured based on the APK. Corresponding chromosomes are distributed according to dot plot order.

**Supplementary figure 6.** Schematic representation of the most conserved GBs associations of *A. edgeworthii* (*Ae*) karyotype as inferred from comparison with the APK. Despite extensive genome reshuffling, the main GB associations involved in the formation of each *Ae* chromosome are indicated by dotted lines in the corresponding APK chromosome colours.

## REFERENCES

Bertioli DJ, Moretzsohn MC, Madsen LH, et al (2009) An analysis of synteny of *Arachis* with *Lotus* and *Medicago* sheds new light on the structure, stability and evolution of legume genomes. BMC Genomics 10:45. https://doi.org/10.1186/1471-2164-10-45

Bonifácio EM, Fonsêca A, Almeida C, et al (2012) Comparative cytogenetic mapping between the lima bean (*Phaseolus lunatus* L.) and the common bean (*P. vulgaris* L.). Theor Appl Genet 124:1513–1520. https://doi.org/10.1007/s00122-012-1806-x

Carvalho CR, Saraiva LS (1993) An Air Drying Technique for Maize Chromosomes without Enzymatic Maceration. Biotechnic & Histochemistry 68:142–145. https://doi.org/10.3109/10520299309104684

Chen F, Dong W, Zhang J, et al (2018) The Sequenced Angiosperm Genomes and Genome Databases. Front Plant Sci 9:418. https://doi.org/10.3389/fpls.2018.00418

Cheng F, Wu J, Wang X (2014) Genome triplication drove the diversification of *Brassica* plants. Horticulture Research 1:1–8. https://doi.org/10.1038/hortres.2014.24

de Oliveira Bustamante F, do Nascimento TH, Montenegro C, et al (2021) Oligo-FISH barcode in beans: a new chromosome identification system. Theor Appl Genet 134:3675–3686. https://doi.org/10.1007/s00122-021-03921-z

do Vale Martins, de Oliveira Bustamante F, da Silva Oliveira AR, et al (2021) BAC- and oligo-FISH mapping reveals chromosome evolution among *Vigna angularis*, *V. unguiculata*, and *Phaseolus vulgaris*. Chromosoma 130(2-3):133–147. https://doi.org/10.1007/s00412-021-00758-9

Faria R, Navarro A (2010) Chromosomal speciation revisited: rearranging theory with pieces of evidence. Trends in Ecology & Evolution 25:660–669. https://doi.org/10.1016/j.tree.2010.07.008

Ferraz ME, Fonsêca A, Pedrosa-Harand A (2020) Multiple and independent rearrangements revealed by comparative cytogenetic mapping in the dysploid Leptostachyus group (*Phaseolus* L., Leguminosae). Chromosome Res 28(3-4):395–405. https://doi.org/10.1007/s10577-020-09644-z

Feulner P, De-Kayne R (2017) Genome evolution, structural rearrangements and speciation. Journal of Evolutionary Biology 30:1488–1490. https://doi.org/10.1111/jeb.13101

Fonsêca A, Ferraz ME, Pedrosa-Harand A (2016) Speeding up chromosome evolution in *Phaseolus:* multiple rearrangements associated with a one-step descending dysploidy. Chromosoma 125:413–421. https://doi.org/10.1007/s00412-015-0548-3

Fonsêca A, Pedrosa-Harand A (2013) Karyotype stability in the genus *Phaseolus* evidenced by the comparative mapping of the wild species *Phaseolus microcarpus*. Genome 56:9. https://doi.org/10.1139/gen-2013-0025

Fuller ZL, Leonard CJ, Young RE, et al (2018) Ancestral polymorphisms explain the role of chromosomal inversions in speciation. PLOS Genetics 14:e1007526. https://doi.org/10.1371/journal.pgen.1007526

Garcia T, Duitama J, Zullo SS, et al (2021) Comprehensive genomic resources related to domestication and crop improvement traits in Lima bean. Nature Communications 12:702. https://doi.org/10.1038/s41467-021-20921-1

Geiser C, Mandáková T, Arrigo N, et al (2016) Repeated Whole-Genome Duplication, Karyotype Reshuffling, and Biased Retention of Stress-Responding Genes in Buckler Mustard. The Plant Cell 28:17–27. https://doi.org/10.1105/tpc.15.00791

Gong Z, Wu Y, Koblízková A, et al (2012) Repeatless and repeat-based centromeres in potato: implications for centromere evolution. Plant Cell 24:3559–3574. https://doi.org/10.1105/tpc.112.100511

Han Y, Zhang T, Thammapichai P, et al (2015) Chromosome-Specific Painting in *Cucumis* Species Using Bulked Oligonucleotides. Genetics 200:771–779. https://doi.org/10.1534/genetics.115.177642

Ho WK, Chai HH, Kendabie P, et al (2017) Integrating genetic maps in bambara groundnut [ *Vigna subterranea* (L) Verdc.] and their syntenic relationships among closely related legumes. BMC Genomics 18:192. https://doi.org/10.1186/s12864-016-3393-8

Hu Q, Ma Y, Mandáková T, et al (2021a) Genome evolution of the psammophyte *Pugionium* for desert adaptation and further speciation. Proc Natl Acad Sci USA 118:e2025711118. https://doi.org/10.1073/pnas.2025711118

Hu T, Chitnis N, Monos D, Dinh A (2021b) Next-generation sequencing technologies: An overview. Human Immunology 82:801–811. https://doi.org/10.1016/j.humimm.2021.02.012

Hufnagel B, Marques A, Soriano A, et al (2020) High-quality genome sequence of white lupin provides insight into soil exploration and seed quality. Nat Commun 11:492. https://doi.org/10.1038/s41467-019-14197-9

Iwata A, Tek AL, Richard MMS, et al (2013) Identification and characterization of functional centromeres of the common bean. Plant J 76(1):47–60. https://doi.org/10.1111/tpj.12269

Iwata-Otsubo A, Lin J-Y, Gill N, Jackson SA (2016) Highly distinct chromosomal structures in cowpea (*Vigna unguiculata*), as revealed by molecular cytogenetic analysis. Chromosome Res 24:197–216. https://doi.org/10.1007/s10577-015-9515-3

Jiao Y, Wickett NJ, Ayyampalayam S, et al (2011) Ancestral polyploidy in seed plants and angiosperms. Nature 473:97–100. https://doi.org/10.1038/nature09916

Kamphuis LG, Williams AH, D’Souza NK, et al (2007) The *Medicago truncatula* reference accession A17 has an aberrant chromosomal configuration. New Phytologist 174:299–303. https://doi.org/10.1111/j.1469-8137.2007.02039.x

Kreplak J, Madoui M-A, Cápal P, et al (2019) A reference genome for pea provides insight into legume genome evolution. Nature Genetics 51:1411–1422. https://doi.org/10.1038/s41588-019-0480-1

Li C, Lin F, An D, et al (2018) Genome Sequencing and Assembly by Long Reads in Plants. Genes 9:6. https://doi.org/10.3390/genes9010006

Li H, Wang W, Lin L, et al (2013) Diversification of the phaseoloid legumes: effects of climate change, range expansion and habit shift. Frontiers in Plant Science 9. https://doi.org/10.3389/fpls.2013.00386

Liao Y, Zhang X, Li B, et al (2018) Comparison of *Oryza sativa* and *Oryza brachyantha* Genomes Reveals Selection-Driven Gene Escape from the Centromeric Regions. Plant Cell 30:1729–1744. https://doi.org/10.1105/tpc.18.00163

Liu Y, Zhang X, Han K, et al (2020) Insights into amphicarpy from the compact genome of the legume *Amphicarpaea edgeworthii*. Plant Biotechnol J pbi.13520. https://doi.org/10.1111/pbi.13520

Lonardi S, Muñoz-Amatriaín M, Liang Q, et al (2019) The genome of cowpea (*Vigna unguiculata* [L.] Walp.). Plant J 98:767–782. https://doi.org/10.1111/tpj.14349

Lyons E, Pedersen B, Kane J, et al (2008) Finding and Comparing Syntenic Regions among *Arabidopsis* and the Outgroups Papaya, Poplar, and Grape: CoGe with Rosids. Plant Physiol 148:1772–1781. https://doi.org/10.1104/pp.108.124867

Lysak MA, Fransz PF, Ali HBM, Schubert I (2002) Chromosome painting in *Arabidopsis thaliana*: Chromosome painting in *Arabidopsis*. The Plant Journal 28:689–697. https://doi.org/10.1046/j.1365-313x.2001.01194.x

Lysak MA, Berr A, Pecinka A, et al (2006) Mechanisms of chromosome number reduction in *Arabidopsis thaliana* and related Brassicaceae species. Proceedings of the National Academy of Sciences 103:5224–5229. https://doi.org/10.1073/pnas.0510791103

Lysak MA, Mandáková T, Schranz ME (2016) Comparative paleogenomics of crucifers: ancestral genomic blocks revisited. Current Opinion in Plant Biology 30:108–115. https://doi.org/10.1016/j.pbi.2016.02.001

Mandáková T, Lysak MA (2018) Post-polyploid diploidization and diversification through dysploid changes. Current Opinion in Plant Biology 42:55–65. https://doi.org/10.1016/j.pbi.2018.03.001

Mandáková T, Hloušková P, Koch MA, Lysak MA (2020) Genome Evolution in Arabideae Was Marked by Frequent Centromere Repositioning. Plant Cell 32:650–665. https://doi.org/10.1105/tpc.19.00557

Martin G, Baurens F-C, Hervouet C, et al (2020) Chromosome reciprocal translocations have accompanied subspecies evolution in bananas. The Plant Journal 104:1698–1711. https://doi.org/10.1111/tpj.15031

McConnell M, Mamidi S, Lee R, et al (2010) Syntenic relationships among legumes revealed using a gene-based genetic linkage map of common bean (*Phaseolus vulgaris* L.). Theor Appl Genet 121:1103–1116. https://doi.org/10.1007/s00122-010-1375-9

Murat F, Armero A, Pont C, et al (2017) Reconstructing the genome of the most recent common ancestor of flowering plants. Nature Genetics 49:490–496. https://doi.org/10.1038/ng.3813

Murat F, Xu J-H, Tannier E, et al (2010) Ancestral grass karyotype reconstruction unravels new mechanisms of genome shuffling as a source of plant evolution. Genome Res 20:1545–1557. https://doi.org/10.1101/gr.109744.110

Noor MAF, Grams KL, Bertucci LA, Reiland J (2001) Chromosomal inversions and the reproductive isolation of species. Proceedings of the National Academy of Sciences 98:12084–12088. https://doi.org/10.1073/pnas.221274498

Oliveira AR da S, Martins L do V, Bustamante F de O, et al (2020) Breaks of macrosynteny and collinearity among moth bean (*Vigna aconitifolia*), cowpea (*V. unguiculata*), and common bean (*Phaseolus vulgaris*). Chromosome Res. https://doi.org/10.1007/s10577-020-09635-0

Parkin IA, Koh C, Tang H, et al (2014) Transcriptome and methylome profiling reveals relics of genome dominance in the mesopolyploid *Brassica oleracea*. Genome Biol 15:R77. https://doi.org/10.1186/gb-2014-15-6-r77

Pavy N, Pelgas B, Laroche J, et al (2012) A spruce gene map infers ancient plant genome reshuffling and subsequent slow evolution in the gymnosperm lineage leading to extant conifers. BMC Biology 10:84. https://doi.org/10.1186/1741-7007-10-84

Pecrix Y, Staton SE, Sallet E, et al (2018) Whole-genome landscape of *Medicago truncatula* symbiotic genes. Nature Plants 4:1017–1025. https://doi.org/10.1038/s41477-018-0286-7

Pellicer J, Hidalgo O, Dodsworth S, Leitch IJ (2018) Genome Size Diversity and Its Impact on the Evolution of Land Plants. Genes 9:88. https://doi.org/10.3390/genes9020088

Qin S, Wu L, Wei K, et al (2019) A draft genome for *Spatholobus suberectus*. Sci Data 6:113. https://doi.org/10.1038/s41597-019-0110-x

Ren L, Huang W, Cannon SB (2019) Reconstruction of ancestral genome reveals chromosome evolution history for selected legume species. New Phytol 223:2090–2103. https://doi.org/10.1111/nph.15770

Ribeiro T, Dos Santos KGB, Richard MMS, et al (2017) Evolutionary dynamics of satellite DNA repeats from *Phaseolus* beans. Protoplasma 254:791–801. https://doi.org/10.1007/s00709-016-0993-8

Ribeiro T, Vasconcelos E, dos Santos KGB, et al (2020) Diversity of repetitive sequences within compact genomes of *Phaseolus* L. beans and allied genera Cajanus L. and Vigna Savi. Chromosome Res 28:139–153. https://doi.org/10.1007/s10577-019-09618-w

Rice A, Glick L, Abadi S, et al (2015) The Chromosome Counts Database (CCDB) – a community resource of plant chromosome numbers. New Phytol 206:19–26. https://doi.org/10.1111/nph.13191

Rieseberg LH (2001) Chromosomal rearrangements and speciation. Trends in Ecology & Evolution 16:351–358. https://doi.org/10.1016/S0169-5347(01)02187-5

Ruprecht C, Lohaus R, Vanneste K, et al (2017) Revisiting ancestral polyploidy in plants. Science Advances 3:e1603195. https://doi.org/10.1126/sciadv.1603195

Schmutz J, Cannon SB, Schlueter J, et al (2010) Genome sequence of the palaeopolyploid soybean. Nature 463:178–183. https://doi.org/10.1038/nature08670

Schmutz J, McClean PE, Mamidi S, et al (2014) A reference genome for common bean and genome-wide analysis of dual domestications. Nat Genet 46:707–713. https://doi.org/10.1038/ng.3008

Schneider KL, Xie Z, Wolfgruber TK, Presting GG (2016) Inbreeding drives maize centromere evolution. Proc Natl Acad Sci USA 113:E987–E996. https://doi.org/10.1073/pnas.1522008113

Schranz M, Lysak M, Mitchellolds T (2006) The ABC’s of comparative genomics in the Brassicaceae: building blocks of crucifer genomes. Trends in Plant Science 11:535–542. https://doi.org/10.1016/j.tplants.2006.09.002

Schubert I (2018) What is behind “centromere repositioning”? Chromosoma 127:229–234. https://doi.org/10.1007/s00412-018-0672-y

Soltis PS, Marchant DB, Van de Peer Y, Soltis DE (2015) Polyploidy and genome evolution in plants. Current Opinion in Genetics & Development 35:119–125. https://doi.org/10.1016/j.gde.2015.11.003

Song X, Sun P, Yuan J, et al (2021) The celery genome sequence reveals sequential paleo-polyploidizations, karyotype evolution and resistance gene reduction in apiales. Plant Biotechnology Journal 19:731–744. https://doi.org/10.1111/pbi.13499

Talbert PB, Henikoff S (2020) What makes a centromere? Experimental Cell Research 389:111895. https://doi.org/10.1016/j.yexcr.2020.111895

The Arabidopsis Genome Initiative (2000) Analysis of the genome sequence of the flowering plant *Arabidopsis thaliana*. Nature 408:796–815. https://doi.org/10.1038/35048692

Vasconcelos EV, de Andrade Fonsêca AF, Pedrosa-Harand A, et al (2015) Intra- and interchromosomal rearrangements between cowpea *[Vigna unguiculata* (L.) Walp.] and common bean (*Phaseolus vulgaris* L.) revealed by BAC-FISH. Chromosome Res 23:253–266. https://doi.org/10.1007/s10577-014-9464-2

Walden N, Nguyen T-P, Mandáková T, et al (2020) Genomic Blocks in *Aethionema arabicum* Support Arabideae as Next Diverging Clade in Brassicaceae. Front Plant Sci 11:719. https://doi.org/10.3389/fpls.2020.00719

Wang J, Sun P, Li Y, et al (2017) Hierarchically Aligning 10 Legume Genomes Establishes a Family-Level Genomics Platform. Plant Physiol 174:284–300. https://doi.org/10.1104/pp.16.01981

Wang J, Zi H, Wang R, et al (2021a) A high-quality chromosome-scale assembly of the centipedegrass *[Eremochloa ophiuroides* (Munro) Hack.] genome provides insights into chromosomal structural evolution and prostrate growth habit. Hortic Res 8:1–13. https://doi.org/10.1038/s41438-021-00636-6

Wang S, Xiao Y, Zhou Z-W, et al (2021b) High-quality reference genome sequences of two coconut cultivars provide insights into evolution of monocot chromosomes and differentiation of fiber content and plant height. Genome Biol 22:304. https://doi.org/10.1186/s13059-021-02522-9

Wendel JF, Jackson SA, Meyers BC, Wing RA (2016) Evolution of plant genome architecture. Genome Biology 17:37. https://doi.org/10.1186/s13059-016-0908-1

Willing E-M, Rawat V, Mandáková T, et al (2015) Genome expansion of *Arabis alpina* linked with retrotransposition and reduced symmetric DNA methylation. Nature Plants 1:14023. https://doi.org/10.1038/nplants.2014.23

Wu S, Han B, Jiao Y (2020) Genetic Contribution of Paleopolyploidy to Adaptive Evolution in Angiosperms. Molecular Plant 13:59–71. https://doi.org/10.1016/j.molp.2019.10.012

Xie D, Xu Y, Wang J, et al (2019) The wax gourd genomes offer insights into the genetic diversity and ancestral cucurbit karyotype. Nature Communications 10:5158. https://doi.org/10.1038/s41467-019-13185-3

Yang L, Koo D-H, Li D, et al (2014) Next-generation sequencing, FISH mapping and synteny-based modeling reveal mechanisms of decreasing dysploidy in *Cucumis*. Plant J 77:16–30. https://doi.org/10.1111/tpj.12355

Zhang H, Koblížková A, Wang K, et al (2014) Boom-Bust Turnovers of Megabase-Sized Centromeric DNA in *Solanum* Species: Rapid Evolution of DNA Sequences Associated with Centromeres. The Plant Cell 26:1436–1447. https://doi.org/10.1105/tpc.114.123877

Zhang S-J, Liu L, Yang R, Wang X (2020a) Genome Size Evolution Mediated by Gypsy Retrotransposons in Brassicaceae. Genomics, Proteomics & Bioinformatics 18:321–332. https://doi.org/10.1016/j.gpb.2018.07.009

Zhang Z, Meng F, Sun P, et al (2020b) An updated explanation of ancestral karyotype changes and reconstruction of evolutionary trajectories to form *Camelina sativa* chromosomes. BMC Genomics 21:705. https://doi.org/10.1186/s12864-020-07081-0

Zhao H, Zeng Z, Koo D-H, et al (2017) Recurrent establishment of de novo centromeres in the pericentromeric region of maize chromosome 3. Chromosome Res 25:299–311. https://doi.org/10.1007/s10577-017-9564-x

Zhao Q, Meng Y, Wang P, et al (2021) Reconstruction of ancestral karyotype illuminates chromosome evolution in the genus *Cucumis*. The Plant Journal 107(4):1243–1259. https://doi.org/10.1111/tpj.15381

Zhuang W, Chen H, Yang M, et al (2019) The genome of cultivated peanut provides insight into legume karyotypes, polyploid evolution and crop domestication. Nat Genet 51:865–876. https://doi.org/10.1038/s41588-019-0402-2

