## Supplementary figures and images for "Comparative cytogenomics reveals genome reshuffling and centromere repositioning in the legume tribe Phaseoleae"

### Supplemental Figure 1

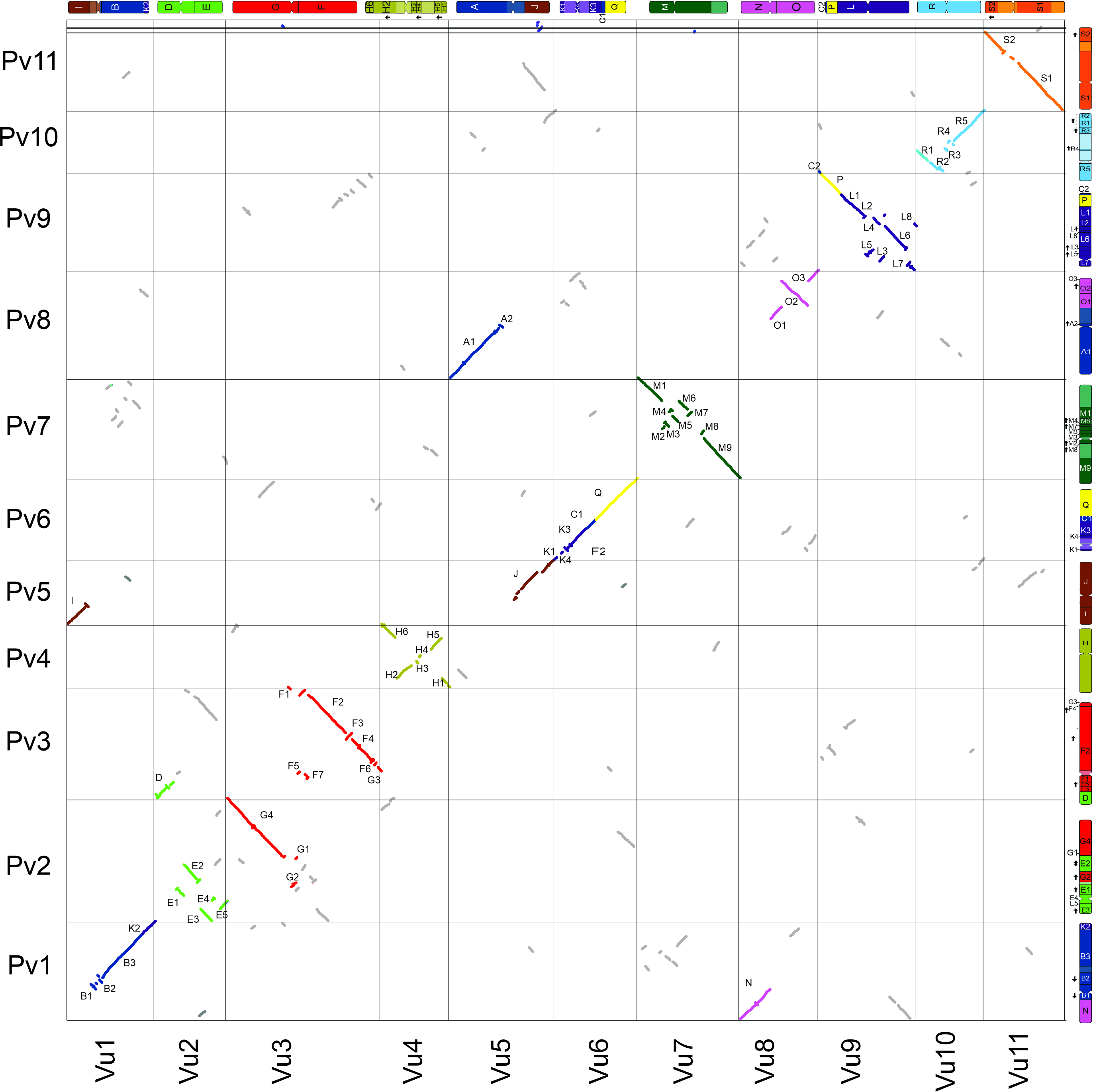

### Supplemental Figure 2

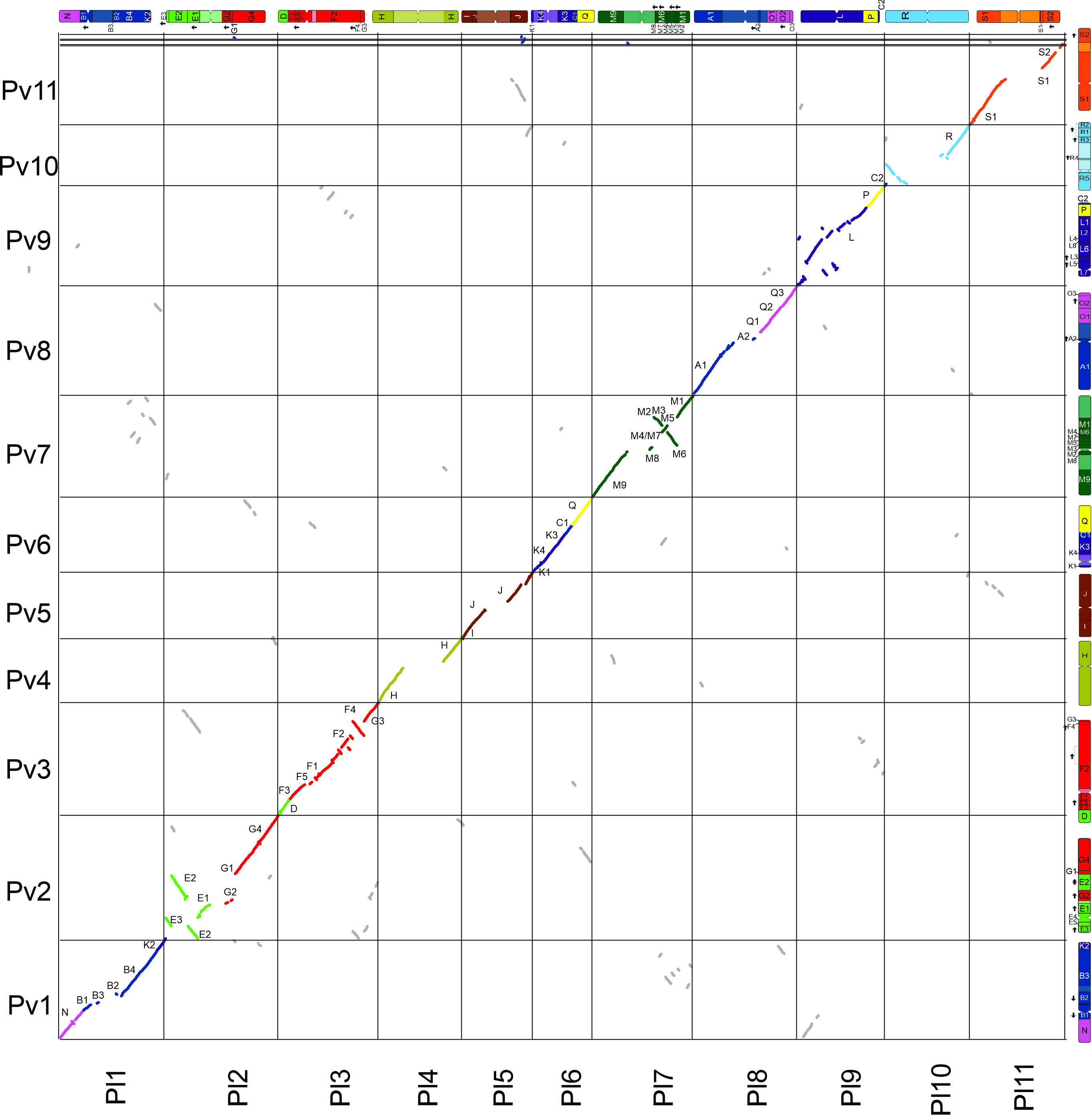

### Supplemental Figure 3

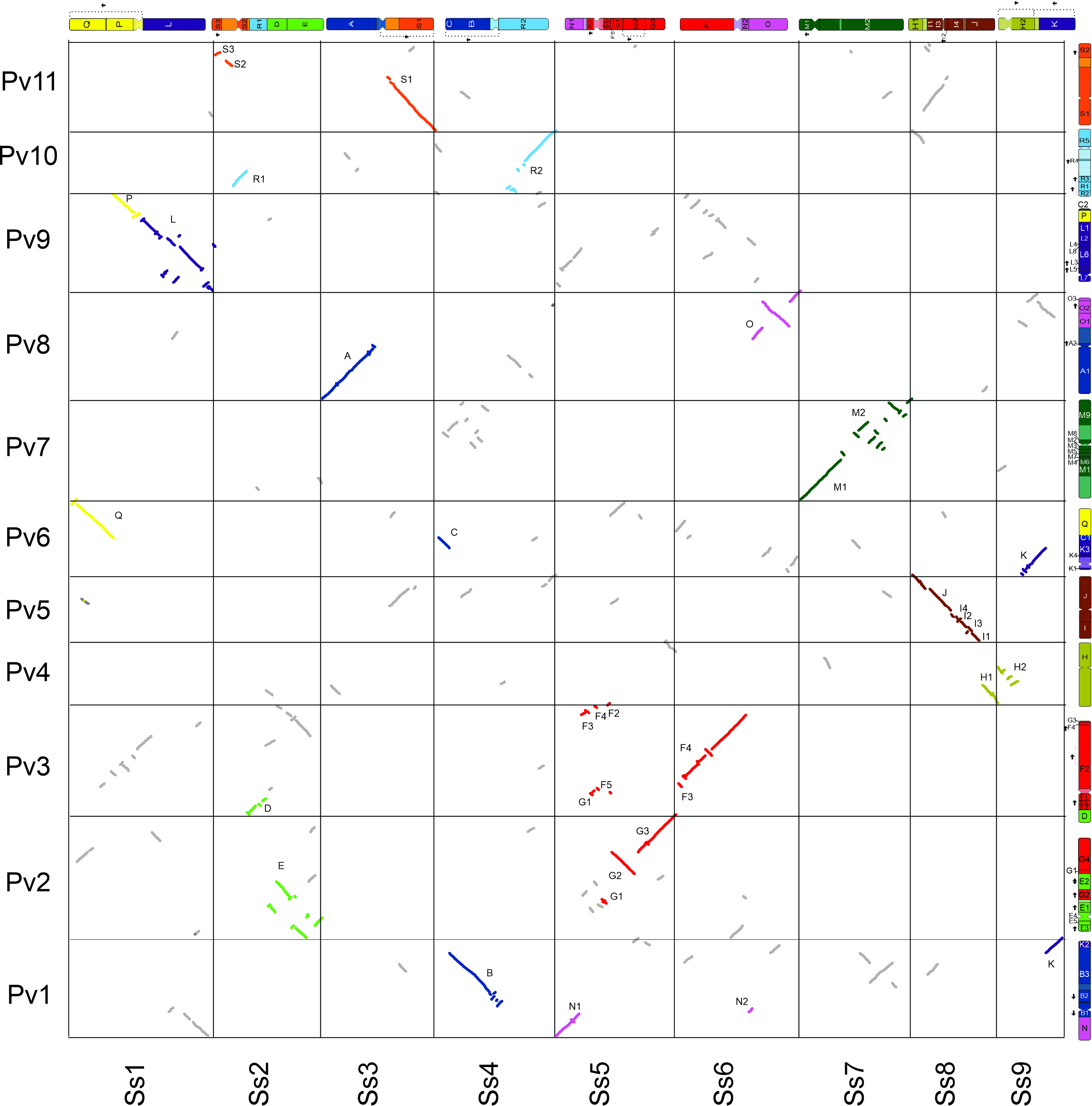

### Supplemental Figure 4

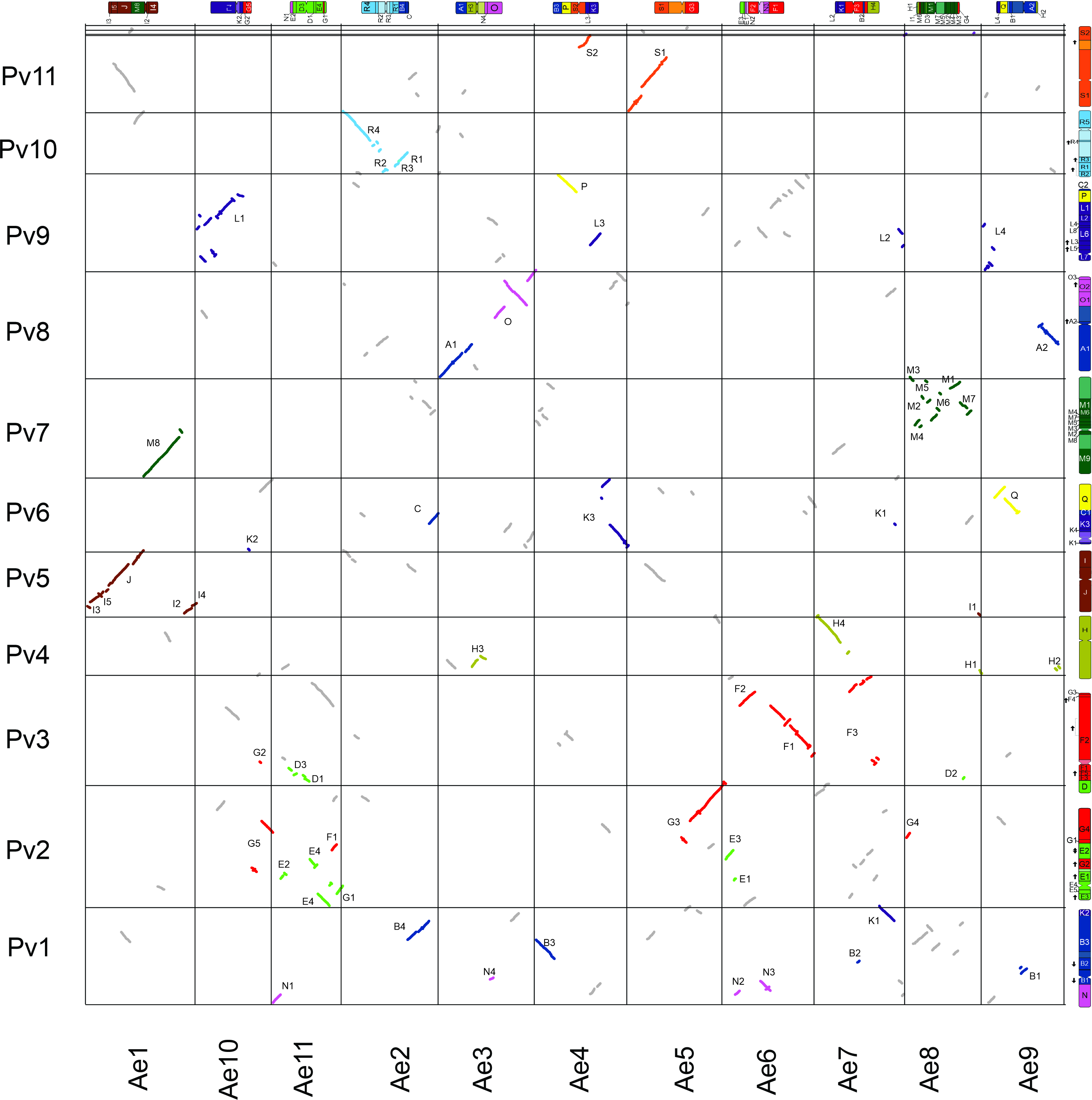

### Supplemental Figure 5

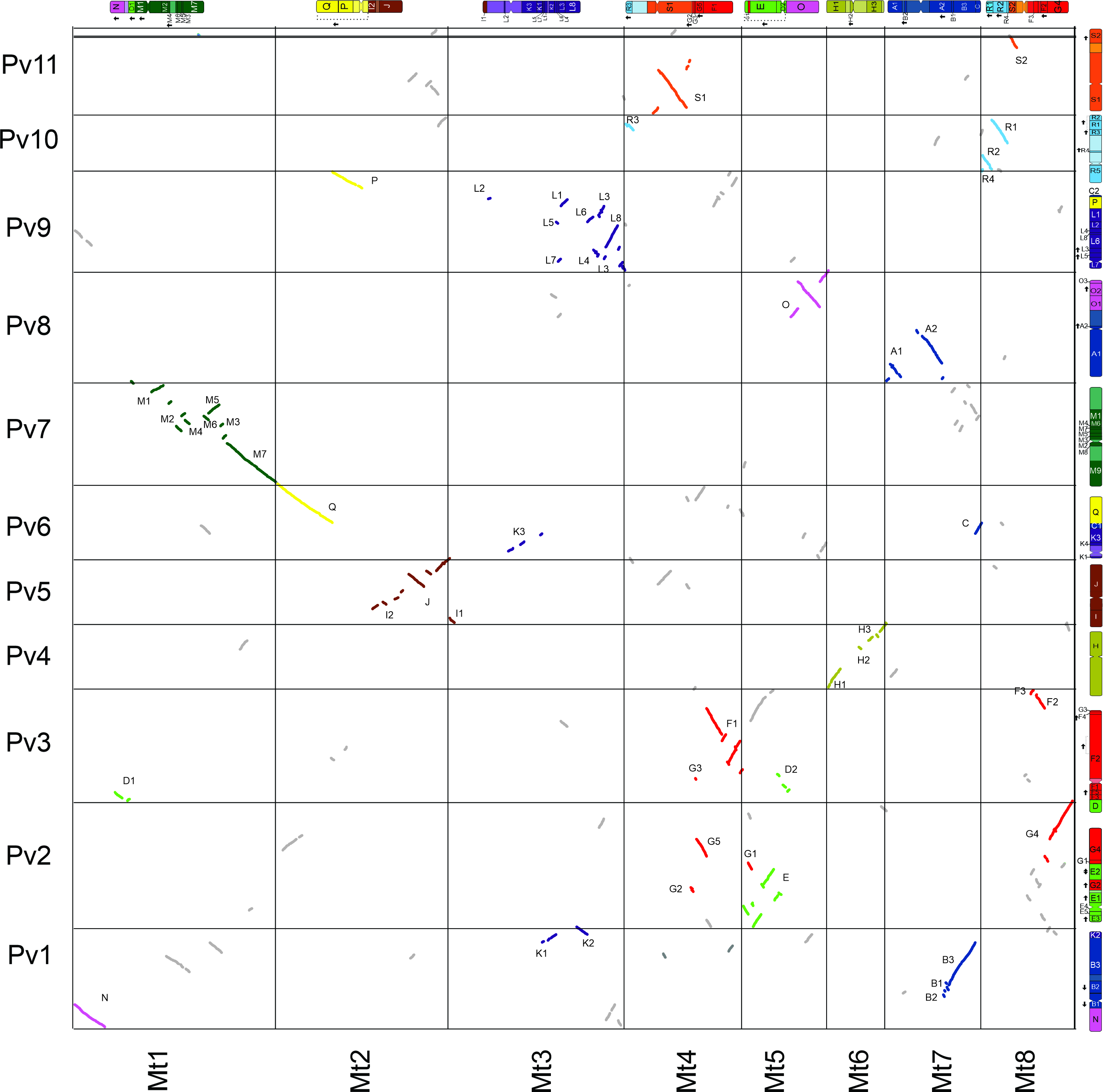

### Supplemental Figure 6

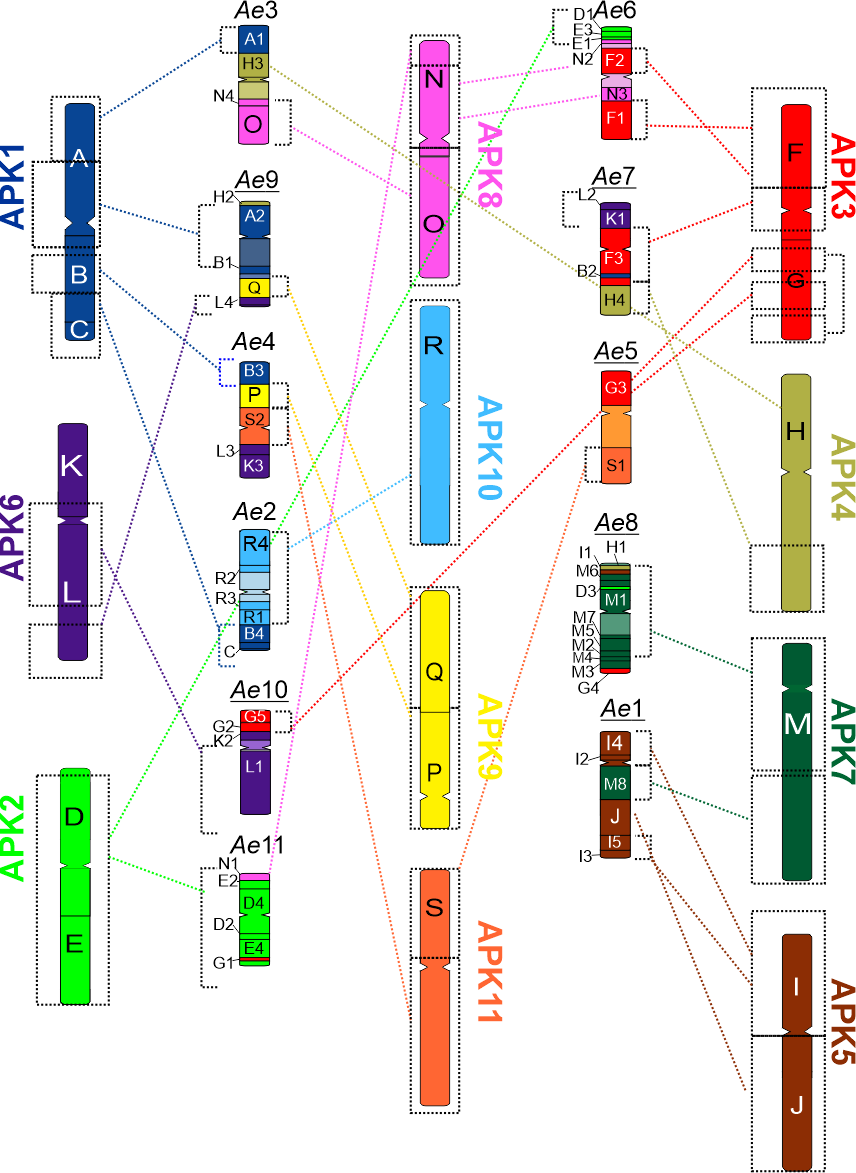
